# A comparison of approaches to scaffolding multiple regions along the 16S rRNA gene for improved resolution

**DOI:** 10.1101/2021.03.23.436606

**Authors:** Justine W Debelius, Michael Robeson, Luisa W. Hugerth, Fredrik Boulund, Weimin Ye, Lars Engstrand

## Abstract

**Motivation:** Full length, high resolution 16s rRNA marker gene sequencing has been challenging historically. Short amplicons provide high accuracy reads with widely available equipment, at the cost of taxonomic resolution. One recent proposal has been to reconstruct multiple amplicons along the full-length marker gene, however no barcode-free computationally tractable approach for this is available. To address this gap, we present Sidle (SMURF Implementation Done to acceLerate Efficiency), an implementation of the Short MUltiple Reads Framework algorithm with a novel tree building approach to reconstruct rRNA genes from individually amplified regions.

**Results:** Using simulated and real data, we compared Sidle to two other approaches of leveraging multiple gene region data. We found that Sidle had the least bias in non-phylogenetic alpha diversity, feature-based measures of beta diversity, and the reconstruction of individual clades. With a curated database, Sidle also provided the most precise species-level resolution.

**Availability and Implementation:** Sidle is available under a BSD 3 license from https://github.com/jwdebelius/q2-sidle

---

Ribosomal RNA marker gene sequencing has been a mainstay of microbiome analysis for more than a decade. While there is a movement toward untargeted metagenomic sequencing, marker gene amplification remains relevant in environments with high host contamination, such as vaginal communities or biopsy samples [1]. However, marker gene sequencing comes with several challenges in taxonomic resolution. The use of short amplicons as opposed to the full 16S rRNA gene has been historically necessary as long read technologies historically had higher error rates and costs than short reads techniques [2–4]. In some cases, these errors exceeded real biological differences between 16S rRNA gene. All amplicon sequencing relies on primers with broad specificity; the primers used to amplify full length 16S genes may not fully capture community diversity [5,6]. However, shorter read technologies also potentially come with drawbacks. Shorter reads from more universal primer pairs may have lower taxonomic resolution than full length sequences and therefore miss important genus- or species-level differences in organisms [7]. Alternatively, organism-specific primers can come at the cost of accurately describing the rest of the community [8,9].

Synthetic long read technology, such as the approach marketed by Loop Genomics, provides the read quality of short read technology with the resolution of long read approaches. Here, short fragments along the full-length marker gene are tagged with a unique molecular identifier before PCR. This approach leverages the lower error rates of short read sequencers coupled with a mostly database-free approach to assembly [10]. However, the technique still uses primers for the full-length sequence, which may not be able to amplify all taxa with equal fidelity. The technique requires full length, unfragmented 16S molecules to work properly, a potential problem for sample types where the DNA may have degraded during storage, like FPVE biopsies embedded biopsies, or sample types which require heavy bead beating, although the specific technique has not been fully benchmarked under these conditions [11,12]

Full length sequences can also be reassemble by amplifying multiple regions along a full length marker gene and then scaffolding using a database approach [6]. This technique may be more robust to random breakage in the DNA. The mix of primers may allow for less overall primer bias. The problem then becomes how to combine the regions. One proposed solution is the use of operational taxonomic units (OTU), clustered against a reference database [13]. A second, user-proposed pipeline relies on regional denoising to generate amplicon sequence variants (ASVs). Taxonomic assignments are made using a naïve Bayesian classifier, and ASVs are scaffolded together using fragment insertion into a reference tree and then profiled using phylogeny-aware metrics. The third potential solution to the problem is the use of the Short Multiple Reads Framework (SMURF) algorithm, which performs regional kmer-based alignment to a reference and then solves the relative abundance using a maximum likelihood estimator model [6]. This allows the use of disjoint regions along a molecular target, and theoretically could be extended to combine multiple marker genes, independent of genome location. The original paper does not consider phylogeny, potentially limiting insights into the microbial community [14]. Additionally, the original implementation was challenging to use and required proprietary software. As a consequence, while the paper has been well cited, the method has not been widely adopted.

To address the issue of combining information from multiple primer regions, we re-implemented the SMURF algorithm and developed a tree building approach, which is released as the q2-sidle (SMURF Implementation Done to acceLerate Efficiency) plugin. Three proposed approaches (closed reference OTUs, ASVs with an insertion tree, and Sidle) were benchmarked to identify the best method for reconstruction, reliably capturing as much sequence information available across multiple gene regions. We further benchmarked the ability of each of these approaches on previously published vaginal microbiome data to determine their ability to recover species-level resolution.

## Materials and Methods

### Implementation

To facilitate reassembly from multiple marker gene regions, we re-implemented the core SMURF algorithm in python as Sidle. The code has 95% test coverage with unit testing. Sidle has been released as a QIIME 2 plugin. This builds on the architecture of the popular microbiome analysis platform, including the decentralized provenance tracking; multiple installation options for Linux, OSX, and virtual boxes for windows operating system; and multiple APIs [15]. Integration with QIIME 2 also creates flexibility: users can enter with fully multiplexed sequences, partially demultiplexed sequences or even a feature table and can select denoising and quality filtering algorithms more appropriate to their data rather than assuming a single quality-filtering error model. To improve performance, Sidle leverages the python Dask distributed computational library for certain pleasantly parallelizable steps in the reconstruction algorithm. Dask allows end users to customize their parallel processing to their local compute architecture, and scales from a single machine to HPC clusters [16].

The Sidle implementation involves five steps: database preparation, regional sample preparation and alignment, table reconstruction, taxonomic annotation, and optionally, building a phylogenetic tree (Figure 1; Supplemental Methods).

**Figure 1.**
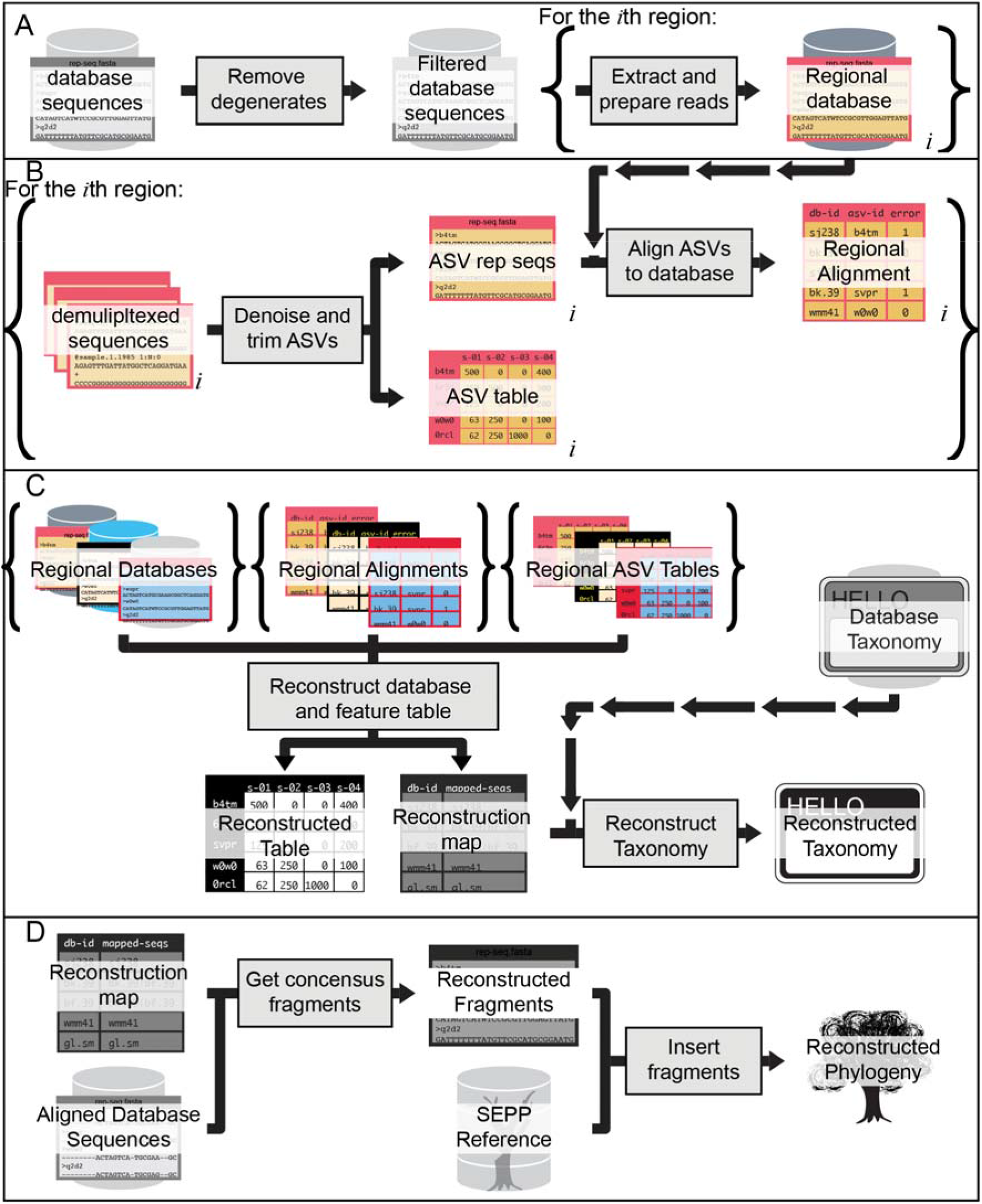
Schematic of Sidle Reconstruction. (A) The database is filtered to remove undesirable sequences and then per-region database are extracted and prepared. (B) The reads for each region are denoised and aligned with the per-region database.(C) The regional databases, regional alignments, and regional ASV tables are combined to reconstruct the database for full length sequences and the feature table. The database map is used with the taxonomy to reconstruct the taxonomy sequences. (D) Optionally, a phylogenetic tree can be reconstructed using the aligned sequences from the reference database to reconstruct fragments, which are inserted into a reference backbone.

### Data Sources

#### Benchmarking data

To benchmark all techniques, we used tutorial dataset provided by the original SMURF paper [6] (Supplemental Methods). This consisted of a single sample without metadata. The sample was compared against the Greengenes 13_5 (SMURF) and Greengenes 13_8 (Sidle) [17].

#### Simulation

We generated a set of reference samples based on previously published experimental data. This provided a base truth community with characteristics similar to true microbiome data and a biologically relevant, if somewhat large, effect size. Amplicons were simulated using *in silico* PCR for three primer pairs (Table S1; Supplemental Methods). Simulations were compared against the Silva 128 database at 99% identity [18].

#### Real Data

We used a set of 24 vaginal samples (8 individuals with 3 replicates) which have been previously described [19] (Table S1; Supplemental Methods). The vaginal samples were compared to the Optivag 16S rRNA database (v0.1) [19]. This curated, vagina-specific database provides accurate species level assignments.

### Reconstruction Methods

All reconstruction methods were performed using the 2020.11 release of QIIME 2 with the Sidle and the RESCRIPt plugins.

#### Closed Reference OTU Clustering

In constructing OTUs, we assumed that denoising had already been applied. Sequences were clustered at 99% identity against the respective reference databases using vsearch (q2-vserach) [20]. Taxonomic assignments were inherited from the database; for the Silva 128 database, the phylogenetic tree was also inherited from the database (Figure S1).

#### ASVs

The feature tables, and their corresponding sequence files, from all regions were merged. Taxonomic classification on the multi region data was performed using a naïve Bayesian classifier trained on the full 16S gene q2-feature-classifer [21]. The final feature table was filtered to exclude any feature without at least phylum level resolution. In cases where the database was unable to classify a taxonomic level, the lowest defined taxonomic level was inherited. For the simulated data, we constructed a phylogenetic tree using fragment insertion into the Silva 128 backbone (q2-fragment-insertion) [22].

#### Multiple region alignment with Sidle

The sidle reference databases were filtered to exclude reference sequences with more than 5 degenerate nucleotides or references which belonged to kingdom Eukaryota. Reference and ASV sequences were trimmed to a consistent length (Table S1). Alignment was performed on a per-region basis allowing no more than 2 nucleotides difference for reads over 300nt and 1 for reads under 300nt. Feature tables were reconstructed using the default parameters in QIIME. Taxonomy was reconstructed, treating missing taxonomic levels as unique designations. For the simulation, the phylogenetic tree was generated using the Silva 128 reference [22,23].

### Performance

We benchmarked the performance of the original SMURF implementation and Sidle on the SMURF tutorial data and the vaginal real dataset (Supplemental Methods). We were unable to run the SMURF code to expand and prepare the database due to a missing function. To profile vaginal samples, we were required to concatenate the files into a single fastq file for each sample and arrange them manually into file folders; this was only possible because the per-region primers had not been trimmed. SMURF was run in MATLAB 2020b (Mathworks, Natick, MA, USA) and profiled with the *profile* function. Sidle was profiled using Snakemake (v 5.3) [24].

### Statistical Analysis of simulated data

Diversity analyses were performed using multiple rarefaction. Feature tables were rarefied to 10,000 sequences/sample five times for each rarefaction method.

#### Alpha diversity

Alpha diversity was characterized using Faith’s Phylogenetic diversity, Observed ASVs, Shannon diversity, and Pielou’s evenness were calculated on each table [25,26]. The relative effect size for alpha diversity metrics was calculated as the absolute value of the Cohen’s d statistic; the mean and standard deviation reflect the five iterations. The values were compared against the reference dataset using an ordinary least squares regression comparing all the iterations for the reconstruction against all reference iterations; reference variation was compared using pairwise testing. Regression was performed using statsmodels (v 0.11.1) [27].

#### Beta Diversity

The effect of reconstruction method on the overall community structure was compared using beta diversity. Rarefied tables were used to calculate Bray-Curtis distance on feature level data; Bray-Curtis distance on a table collapsed to genus level; weighted UniFrac; and unweighted UniFrac distances (q2-diversity) [28–30]. The reconstruction methods were compared to the reference dataset using Mantel’s test with 999 permutations in scikit-bio (v. 0.5.5; www.scikit-bio.org) [31]. The correlation presented is the average Mantel correlation across all pairwise mantel tests. The mantel correlation for reconstruction methods were compared using a one-sided t-test with unequal variance. The t-test was calculated in scipy (v 1.5.2) [32].

#### Clade Abundance

Taxonomic abundance was compared between organisms using an ordinary least-squares regression. Since the reference data and assemblies were annotated using different databases, we first harmonized the taxonomy. To simplify the comparison, organisms in the reconstructed taxonomy (based on the Silva database) which were labeled as “ambiguous”, “unidentified” or “uncultured” were treated as the equivalent of the un-annotated levels in the Greengenes (reference) database. We also treated any level where the taxonomic classifier could not resolve the organism or where Sidle could not resolve the taxa as unannotated. Unannotated levels inherited the lowest defined level. Class assignments were harmonized between the reference and reconstruction database.

The counts were normalized and filtered to retain features at the specified level that were present with an average abundance of at least 0.01%. We used linear regression with a zero-intercept to calculate the correlation between the reference and reconstructed taxonomic abundance. We evaluated the relationship between the values using a paired t-test with a Bonferroni corrected p-value. We considered a p < 0.05 with at least 5% deviation to be significant. Modeling was performed in statsmodels and scipy [27,32]. The ratio between the reconstruction and reference abundance was plotted using seaborn (v. 0.11.0) and matplotlib (v. 3.2.2) [33,34].

### Statistical Analysis of Real Data

Within-subject stability was calculated using Bray-Curtis distance on the species-level data for each method. We used a linear mixed effects model using each individual as a random effect; modeling was performed in statsmodels [27]. We calculated the distance from the single region sample by filtering the data to retain species present with a relative abundant of at least 10% in at least one pool (n=11); these features represented at least 89% of the relative abundance for all 8 pools. In cases where assignments were different (i.e. cases where the level could not be assigned or resolved), the missing values were treated as 0 counts. A PCoA projection and corresponding biplot was calculated using q2-diversity; the PCoA was visualized using q2-Emperor [35].

## Results

### A comparison of Sidle and SMURF

We first compared the performance of Sidle and SMURF for database preparation, profiling a single sample, and profiling multiple samples (Table S2). We first tried to prepare the Greengenes database using the set of six SMURF primers [6,17]. We were unable to profile the full SMURF database generation because the SMURF library was missing a necessary function to expand the degenerate sequences. This also prevents the use of other databases directly through the SMURF implementation. Even with this function excluded, database processing with SMURF took an hour and 43 minutes, compared to the 27 minutes required by Sidle, a three-fold increase (Table S2). SMURF was more efficient at single sample profiling, taking 7:56 compared to Sidle’s 35:46; this was primarily due to differences in the time spent on denoising and reconstruction. The Sidle implementation “solves” the database on the fly, determining the correct sequences during reconstruction while SMURF determines this database structure during the database preparation step.

We then tried profiling the two functions using a real dataset rather than the provided tutorial data. We first tried using a curated, environment-specific database with SMURF, however, due to the missing function, we were unable to prepare this database. We therefore used the pre-expanded Greengenes database with the two regional primers. It took SMURF 88 minutes to prepare the database, and when we tried to use this database with the samples, the database mapping was incorrect and data could not be processed. In contrast, it took 15 minutes to prepare the Greengenes database with Sidle, and full reconstruction took less than 30 minutes, for a total run time of 44 minutes for 24 samples.

Having determined that Sidle was a more runnable implementation, we then explored the effect of different reconstruction methods on the reconstructed community. We tried three methods: using closed reference OTU clustering (“OTUs”), ASVs with naïve Bayesian taxonomic assignment and a fragment insertion tree (“ASVs”) and multiple region reconstruction (“Sidle”). These were performed starting from the same set of simulated amplicons.

### Community Structure

We found the reconstructed alpha diversity was highly correlated with the reference values (R^2^ > 0.85, Table 1). All three reconstruction methods over-estimated the phylogenetic diversity, although the over-estimation was greater when fragment insertion was used, resulting in 2.91 fold over estimation with Sidle and 3.53 fold over-estimation using ASVs for reconstruction. With non-phylogenetic metrics, Sidle most faithfully reconstructed the alpha diversity with no over-estimation, within 0.1 fold (Table 1). ASVs consistently over-estimated the alpha diversity metrics by the largest factor. OTUs fell between, overestimating compared to the reference and sidle.

**Table 1.** The effect of reconstruction method on the observed alpha diversity

| **Reconstruction**<br>**Method** | **Biological effect** |  | **Change from reference** |  |  |
| --- | --- | --- | --- | --- | --- |
|  | **mean** | **(std)** | **mean** | **(std)** | **R<sup>2</sup>** |
| Phylogenetic Diversity |  |  |  |  |  |
| Reference<sup>a</sup> | 3.92 | (0.04) | 1.000 | (0.002) | 0.995 |
| OTUs | 4.13 | (0.07) | 1.309 | (0.003) | 0.993 |
| ASVs | 4.23 | (0.15) | 3.523 | (0.007) | 0.992 |
| Sidle | 4.19 | (0.11) | 2.910 | (0.009) | 0.992 |
| Observed Features |  |  |  |  |  |
| Reference<sup>a</sup> | 5.03 | (0.05) | 1.000 | (0.002) | 0.996 |
| OTUs | 5.02 | (0.10) | 1.337 | (0.003) | 0.995 |
| ASVs | 4.89 | (0.12) | 1.869 | (0.006) | 0.995 |
| Sidle | 4.84 | (0.11) | 0.998 | (0.003) | 0.995 |
| Shannon Diversity |  |  |  |  |  |
| Reference<sup>a</sup> | 4.4 | (0.04) | 1.000 | (0.000) | 0.999 |
| OTUs | 4.21 | (0.04) | 1.102 | (0.000) | 0.997 |
| ASVs | 4.41 | (0.05) | 1.188 | (0.001) | 0.998 |
| Sidle | 4.32 | (0.06) | 1.000 | (0.001) | 0.998 |
| Pielou's Evenness |  |  |  |  |  |
| Reference<sup>a</sup> | 3.59 | (0.07) | 1.000 | (0.000) | 0.998 |
| OTUs | 3.28 | (0.05) | 1.053 | (0.000) | 0.990 |
| ASVs | 3.66 | (0.06) | 1.078 | (0.000) | 0.995 |
| Sidle | 3.52 | (0.02) | 1.001 | (0.001) | 0.994 |

<sup>a</sup>Reference is compared to itself through multiple rarefaction

We also explored beta diversity (Table 2). We found a strong correlation between the reference community and the reconstructed community using all three reconstruction methods across all four metrics (mantel R^2^ > 0.90, p=0.001, 999 permutations). However, Sidle represented a significant improvement over OTU clustering and ASV reconstruction for unweighted UniFrac (p < 0.002), weighted UniFrac (p < 1×10^−12^), and feature-based Bray-Curtis distance (p < 1×10^−12^). However, it underperformed on genus-level Bray Curtis distance, where ASV-based analysis performed best (p < 1×10^−8^).

**Table 2.** The effect of reconstruction method on the observed beta diversity

| Method | Biological effect size |  |  | Comparison to reference |  |  |
| --- | --- | --- | --- | --- | --- | --- |
|  | Mean | R<sup>2</sup><br>(std) | p-value<sup>a</sup> | mean | R<sup>2</sup><br>(std) | p-value<sup>a</sup> |
| Uweighted UniFrac |  |  |  |  |  |  |
| Reference<sup>b</sup> | 0.677 | (0.007) | 0.001 | 0.984 | (0.001) | 0.001 |
| OTUs | 0.669 | (0.011) | 0.001 | 0.979 | (0.002) | 0.001 |
| ASVs | 0.618 | (0.012) | 0.001 | 0.976 | (0.002) | 0.001 |
| Sidle | 0.679 | (0.010) | 0.001 | 0.980 | (0.002) | 0.001 |
| Weighted UniFrac |  |  |  |  |  |  |
| Reference<sup>b</sup> | 0.863 | (0.002) | 0.001 | 0.999 | (0.000) | 0.001 |
| OTUs | 0.842 | (0.001) | 0.001 | 0.975 | (0.000) | 0.001 |
| ASVs | 0.826 | (0.001) | 0.001 | 0.974 | (0.000) | 0.001 |
| Sidle | 0.841 | (0.001) | 0.001 | 0.978 | (0.001) | 0.001 |
| Bray Curtis |  |  |  |  |  |  |
| Reference<sup>b</sup> | 0.835 | (0.001) | 0.001 | 0.999 | (0.000) | 0.001 |
| OTUs | 0.826 | (0.001) | 0.001 | 0.998 | (0.000) | 0.001 |
| ASVs | 0.819 | (0.001) | 0.001 | 0.998 | (0.000) | 0.001 |
| Sidle | 0.839 | (0.002) | 0.001 | 0.999 | (0.000) | 0.001 |
| Genus level Bray Curtis |  |  |  |  |  |  |
| Reference<sup>b</sup> | 0.862 | (0.001) | 0.001 | 0.999 | (0.000) | 0.001 |
| OTUs | 0.862 | (0.001) | 0.001 | 0.995 | (0.000) | 0.001 |
| ASVs | 0.860 | (0.000) | 0.001 | 0.995 | (0.000) | 0.001 |
| Sidle | 0.863 | (0.001) | 0.001 | 0.995 | (0.000) | 0.001 |

<sup>a</sup> Permutative p-value with 999 permutations
<sup>b</sup> Reference is compared to itself through multiple rarefaction

### Taxonomy

We compared the correlation between the relative abundance of collapsed taxa at the class level. Database harmonization across the lower taxonomic levels is notoriously difficult, and we found large differences below class level.

We identified a total of 12 classes present with an average relative abundance of at least 0.01% in the reference dataset which could be mapped to the reconstructed data (Figure 2, Table S2). We found 8 classes where at least one reconstruction method was significantly different (p < 0.05; > 5% deviation). This 5% threshold was selected because of compositionality in the data: to have a relative increase in one class, we must lose relative abundance somewhere else; by selecting this threshold, we hoped to allow some shifts associated with compositionality in the data. We found that all three reconstruction methods consistently underestimated class Clostridia (p. Firmicutes) by between 10% and 8% (OTUs 0.92 [95% CI 0.90, 0.93]; ASVs 0.90 [95% CI 0.89, 0.92], Sidle 0.91 [95% CI 0.89, 0.93]). We also found an over-estimation of class Mollicutes (p. Tenericutes). However, while OTU clustering and ASVs over-estimated the relative abundance by 22% [95% CI 19%, 25%] (p < 0.005) for both methods, Sidle only over-estimated by 8% [95% CI 6%, 11%] (p=0.03). We also found Sidle performed better in reconstruction of classes Erysipelotrichia and Deltaproteobacteria, which OTU and ASV reconstruction overestimated and in class Epsilonproteobacteria, which OTU and ASV-based reconstruction significantly underestimated (Table S2). Overall, ASV-based methods had significant deviation from the reference in 8 classes, OTU clustering missed in 5 classes, and Sidle underperformed in two cases.

**Figure 2.**
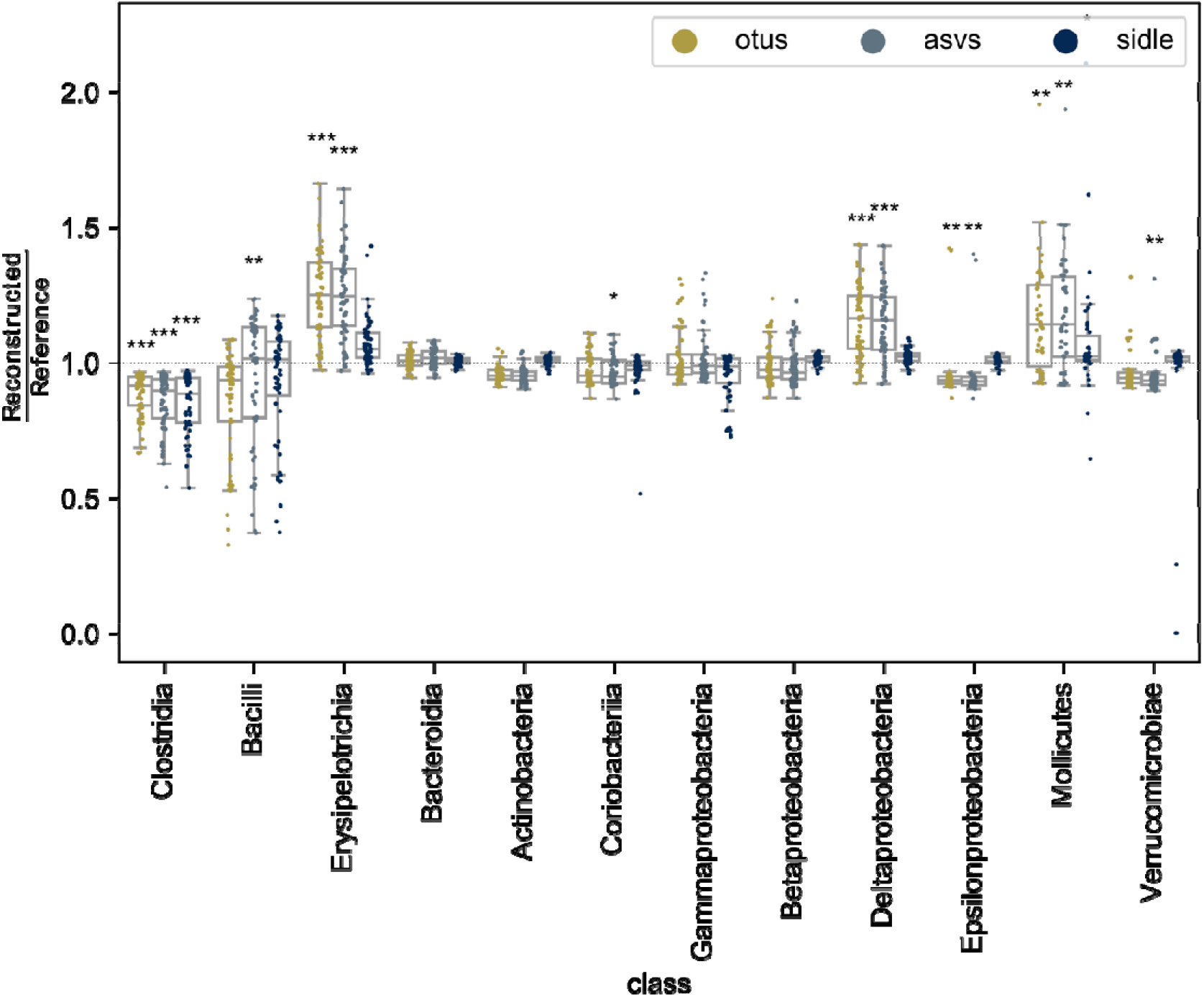
Reconstruction method affects the observed relative abundance of bacterial classes. The ratio of the reconstruction method (OTUs: yellow, ASVs: silver, sidle: dark blue) shows differences in the reconstruction accuracy. The boxplots with at least a 5% deviation on average are labeled with FDR-corrected p-value: * p < 0.05; ** p < 0.01; *** p < 0.001. Values below 1 represent under-estimation of a given clade, while those above 1 represent over-estimation of the given clade.

### Applications to real data

We also explored the effect of reconstruction on real data using a curated, species-level database. We first looked at reconstruction using vaginal samples from eight individuals (Figure 2). We compared a current approach – ASVs drawn from a single region to samples reconstructed using OTUs, ASVs from both V13 and V34 regions, and Sidle annotated with Optivag database. Optivag is an environment specific, manually curated database, designed to allow accurate species level annotation in vaginal communities [19].

We first evaluated the ability of the ASV-based methods and Sidle to resolve taxonomy. With ASVs from the V34 region alone, the naïve Bayesian classifier was unable to resolve species level resolution for a total of 111 ASVs. Classification using the full 16S rRNA gene sequence with both regions led to 321 ASVs unclassified at species level, including 62 ASVs of 192 mapped to genus *Lactobacillus*. Sidle was unable to resolve one feature, which led to one unresolved species: a genus member of *Streptococcus* mapped to either *Streptococcus infantis* or *Streptococcus oralis*.

We next looked at the taxonomic composition of the individuals using species-level data. We found the individual vaginal composition was relatively stable, regardless of the method used. OTUs were significantly more stable than collapsed ASVs from multiple regions (p=0.007); there were not significant differences in stability between any other pairs of metrics (Figure 3A). We also found the individual to be the strongest determinant of the community structure. One major concern was the inability of either ASV-based method to accurately resolve *Lactobacillus* species making it potentially difficult to accurately distinguish between *Lactobacillus crispatus* and other species (Figure 3B,C).

**Figure 3.**
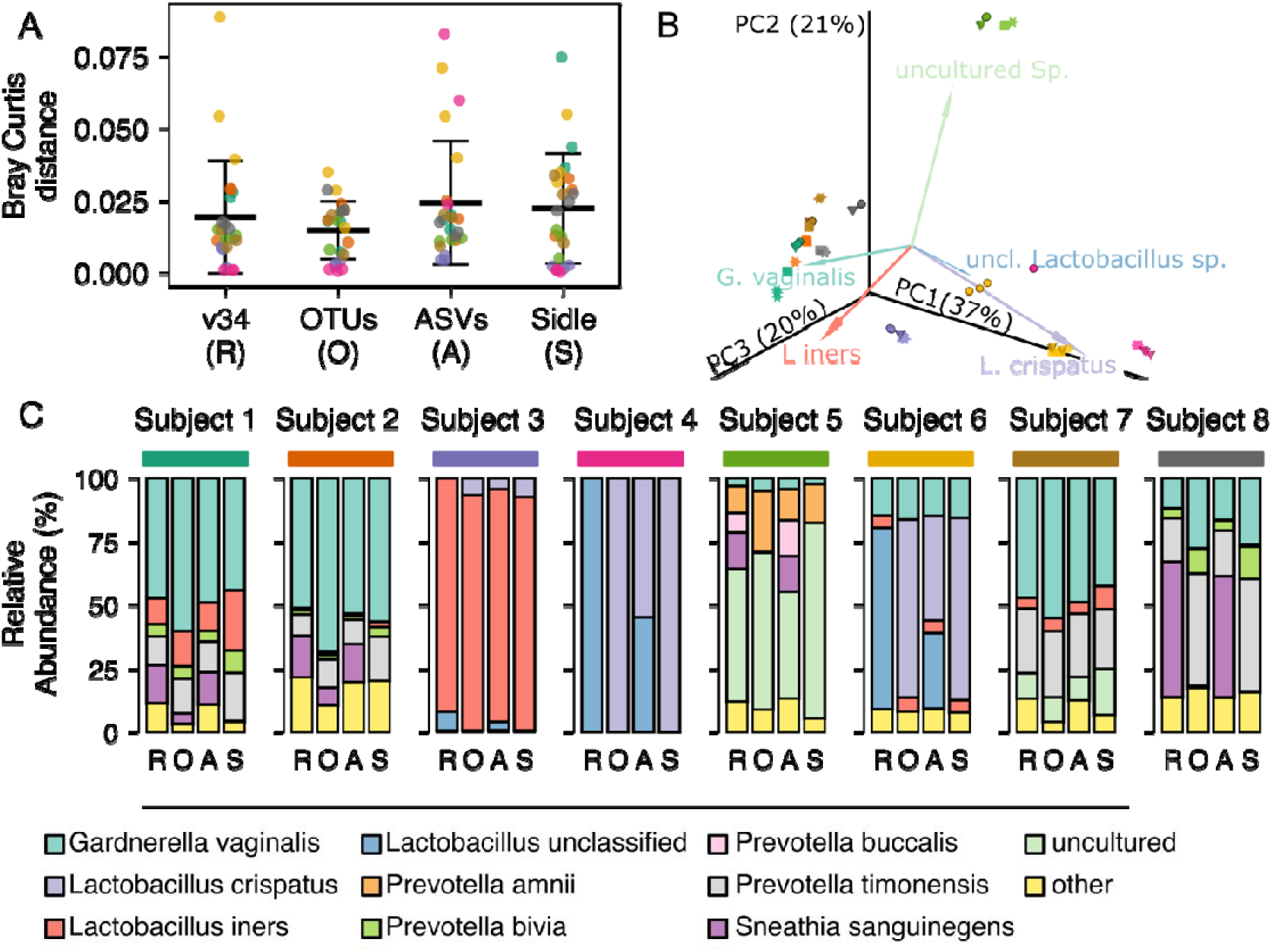
The effect of reconstruction method on vaginal communities at species level resolution. (A) The within pool species level Bray Curtis distance for reconstruction with the V34 region only (R), OTUs (O), ASVs (A), and Sidle (S). Points show intra-subject distance, colored by subject. The black bar indicates the global mean, error bars are the standard deviation. (B) PCoA biplot of Bray Curtis distance on species level data combined across methods. Points are colored by subject (matching A and C), shape indicates the reconstruction methods (circle: v34 only, square: OTUs, star: cone: ASVs; star: Sidle). The five most abundant clades are shown in the biplot. (C) The average relative abundance per subject for each of the four reconstruction methods.

## Discussion

In this analysis, we explored three methods for reconstructing multiple fragments of a larger target gene using a reference database. Our results suggest that the Sidle implementation of the SMURF algorithm was the best method for reconstructing microbial composition from multiple 16S rRNA gene regions. In simulation studies, Sidle most accurately calculated non-phylogenetic alpha diversity, feature-based beta diversity, and led to the lowest bias in clade relative abundance. Interestingly, we found the tree building method was associated with the observed phylogenetic diversity. Both ASV reconstruction and Sidle rely on a fragment-insertion based approach, where the sequences are inserted into a reference backbone [22]. Placements near the tips appear to potentially expand the distance. However, this effect did not extend to community comparisons. However, although the insertion tree affected the phylogenetic alpha diversity, it did not affect the UniFrac distance (beta diversity) between samples, suggesting this may not be a major drawback for such metrics.

Using real data and an environment-specific curated database, we also found that Sidle reconstruction provided the most precise species-level annotation. For vaginal communities like the example community used in this analysis, accurate, species-level *Lactobacillus* assignments are crucial because closely related species have different effects on community structure [36]. For example, one study found that vaginal communities containing *L. crispatus* but not *L. inners* was able to inhibit *E. coli* growth [37]. In our data, the classifier was unable to identify ASVs which were likely *L. crispatus* at the species level, leaving them annotated as an unclassified member of genus *Lactobacillus*. The inability of ASV-based annotation to perform species-level resolution therefore has implications for our biological understanding.

Sidle had overall superior performance: with only two regions of the 16S rRNA gene, we were able to resolve the species for all but one feature, which was annotated to genus level. In contrast, our full-length naïve Bayesian classifier that was used for ASV-based annotation was unable to assign taxonomy for 111 ASVs, including several members of genus *Lactobacillus*. It is possible that if we had combined region-specific classifiers, we might have improved the taxonomic resolution, however, this might also create a bias since different classifiers would be used on different regions [21]. The evaluation is perhaps hardest with OTU clustering. Because the OTUs use the taxonomic annotation assigned to the reference sequence in the database, the observed sequence annotation depends on the database resolution. However, recent work has suggested that the traditional 97% identity threshold used for OTU clustering is insufficient for species-level annotation, and short read amplicons require almost 100% identity OTUs (essentially ASVs) [38]. It has also been argued that reference based OTU clustering methods can be misleading: the sequences included in the OTU clusters may have a similarity larger than the threshold identity, as long as they share the same level of similarity to the reference [39]. The main advantage of OTU clustering for multiple region scaffolding is that the use of consistent reference allows multiple regions to be combined, however, the approach comes with all the drawbacks of single region OTUs-clustering.

Although Sidle performed the best of our reconstruction methods, there are some drawbacks. First, although the authors of the SMURF algorithm claim species level resolution, this is obviously limited by database resolution. With specialized, well curated databases like the Optivag database or Human Oral Microbiome Database, species level resolution is achievable and trust-worthy [19,40,41]. However, more general databases like the Silva database may not provide accurate annotation at lower taxonomic levels, especially because Silva does not curate species assignments [23]. Therefore, the user must consider the database they plan to use and its resolution. Next, Sidle and OTU clustering are limited by database coverage. The methods may not be appropriate for environments with poor database coverage, such as soil or saltwater, since sequences may be discarded. Third, the SMURF algorithm (and Sidle by extension) requires the exact primers used to amplify the sequences for database preparation. Databases are re-usable, so companies with proprietary primers might be able to provide a prepared database. However, this may be a challenge for data re-use and future publications will need to be careful about including primer pairs and read lengths used for annotation.

In conclusion, we present Sidle, an open-source implementation of the SMURF algorithm with a novel tree building approach. We demonstrated that Sidle was best able to reconstruct a reference community in reconstruction and provided high quality species level annotation with a curated database. We hope this library serves as a resource to the community.

## Author Contributions

JWD wrote the q2-sidle plugin; LWH and MR reviewed the code. JWD designed the simulation experiment, performed the simulation, and analyzed the real data. JWD wrote the manuscript with critical edits from LWH and MR. LE and WY secured funding. All authors reviewed and approved the final manuscript.

## Acknowledgements

The authors wish to thank Noam Shental for his permission to re-implement the Matlab SMURF code in python under a BSD-3 license. Thanks to the QIIME 2 forum community for fruitful discussions about how to separate and evaluate multiple 16S regions and invaluable support for plugin development, especially Evan Bolyen, Matthew R Dillion, Carli Jones, and Chris Keefe. Thanks also to Nele Brussellaers for her helpful discussion and edits.

## Data Availability

All work is based on published datasets. Simulations are based on Yatsunenko et al. Original sequences can be downloaded from ENA study PRJEB3079; simulation seed data came from Qiita study 850, using the 100nt 97% closed reference OTU table (Qiita artifact 45113).

Real data is derived from a benchmarking study by Hugerth et al. Sequences are deposited in ENA under study PRJEB37382. Data used from comparing the MATLAB SMURF implementation and Sidle performance came from Fuks et al via their tutorial; data was downloaded from https://github.com/NoamShental/SMURF.

## Code Availability and Implementation

The q2-sidle plugin is available as a pip-installable qiime2 plugin under a BSD 3 license (https://github.com/jwdebelius/q2-sidle). For installation instructions and tutorials, see https://q2-sidle.readthedocs.io/en/latest/.

Analysis code for this paper is available from https://github.com/jwdebelius/avengers-assemble

