## Supplemental File for "A comparison of approaches to scaffolding multiple regions along the 16S rRNA gene for improved resolution"

**Supplemental Material**

Supplementals Methods.

**Tables**

**Table S1.** Regional Primers and trim length

**Table S2.** Runtime comparisons for SMURF and Sidle

**Table S3.** Comparison of reconstruction methods on class level data

**Figures**

**Figure S1.** OTU processing workflow.

**Figure S2**. ASV processing workflow

**Supplemental Methods**

*Details of Sidle Implementation*

Per-region sidle databases. For complex or large databases, the full-length sequences are filtered to exclude databases with excessive degenerate nucleotides using q2-sidle. (The original paper defined this as 3 nucleotides, we have defined this as 5 nucleotides.) This limits computational requirements by decreasing the number of sequences required for alignment. Additionally, taxonomic pre-filtering can be applied to remove any undesired reference sequences (i.e. chloroplasts, mitochondria, under annotated database sequences) through the qiime2 taxonomy plugin (q2-taxa) [1]. The alignment-based algorithm will discard reads associated with an unassigned reference, and therefore specialized environmental databases can be further curated. This curation improves runtime. The local reference database for each region is then constructed by extracting regional reference sequences using the qiime2 feature classifier plugin with default parameters [2]. The regions can then be formatted for Sidle alignment by expanding degenerate sequences and removing duplicates using the q2-sidle plugin.

Per-region feature table preparation. The next step in reconstruction is to prepare the regional tables. Data can enter the pipeline full multiplexed, demultiplexed into samples, demultiplexed into samples and regions, or as amplicon sequence variant (ASV) tables. Sidle relies on the QIIME 2 capabilities to demultiplex the data as appropriate, using either the built in per-sample demultiplexing or through the q2-cutadapt plugin [3]. If the data is multiplexed by region, Cutadapt can also be used to split samples into regions after barcode trimming. The tables can then be denoised using the author’s preferred denoising method. QIIME 2 includes two denoising algorithms: deblur [4] and DADA2 [5]. Deblur was recommended by the authors of the original paper and may have shorter regional runtime than DADA2, however the DADA2 algorithm is more flexible and recent updates can accurately denoise more sequencing types, *e.g.* Ion Torrent or 454 sequencing. If dada2 is used, an optional script in q2-sidle allows the user to trim their ASV representative sequences to a fixed length, since this is required for alignment. Denoising generate a set of representative sequences and a table which maps the number of times the sequence was observed in each sample.

Regional Alignment. The ASV representative sequences are then aligned to the reference database allowing a fixed number of mismatches per region between the reference reads and ASVs. The algorithm allows the use of different read lengths in different regions, and correspondingly, different regional mismatches. Alignment leverages the dask library for parallel computation, since the process is pleasantly parallel [6,7].

Reconstruction. The feature table and taxonomy are reconstructed in two main steps. First, the input database is “solved”, determining the final reconstructed database identifiers based on the regions used during the process. Although this adds computational time during reconstruction, it increases the flexibility since it allows users to mix and match their reference database regions. Once the database is reconstructed, the relative abundance of each sample is solved through the maximum likelihood estimation method presented in the original paper [8]. Although each sample must be solved individually, parallel computing again increases the speed of this step. The final reconstruction creates a scaled feature table with frequency represented in region-weighted counts, and a database map, describing the relationship between the original sequences in the database and their final assignment.

In cases where database sequences were fully resolved, the taxonomic assignments are inherited from the database. In cases where database sequences could not be resolved based on the regions analyzed, taxonomy was inherited to the lowest common ancestor.

Tree building. Sidle introduces a tree build approach for multiple region reconstruction based on fragment insertion into a reference tree [9]. If fragments could be resolved to a single reference sequence, we used the original position in the tree. In cases where multiple reference sequences mapped to the same reconstruction fragments, the representative sequence was determined as the consensus sequence between the most extreme primers. Fragments can then be inserted into the reference tree. However, this phylogenetic reconstruction step requires a fragment insertion tree be available for the database which may limit the ability to perform phylogenetic reconstruction on more specialized databases.

*Benchmark Data*

The tutorial sample was downloaded and imported into QIIME as paired-end sequences. The data was demultiplexed into individual regions and primers were trimmed using q2-cutadapt [3]. Paired end reads were joined using q2-vserach, quality filtered with q2-quality-control, and denoised using q2-deblur using default parameters. All paired end ASVs were trimmed to 125nt, except reads from region 1 (74F-315R) which were trimmed to 215nt and from region 6 (1143F-1336R), which were trimmed to 175nt (Table S1) [4,10]. Alignments allowed a single mismatch.

In MATLAB, reads were split into regions and quality filtered using the default parameters in the SMURF profile_one_sample.m script [8].

*Data Simulation and Processing*

Data from a study of fecal samples from infants and adults across three countries was used as a seed for simulated communities [11]. The 97% closed reference OTU table trimmed to 100nt and sample metadata for fecal samples were downloaded from qiita.ucsd.edu (Study ID 850) [12]. Samples from children younger than 3 years old were categorized as “infant” (n=137), samples from individuals between 20 and 60 were categorized as “adult” (n=171). We mapped the OTUs back to the full-length sequences in the Greengenes database and excluded any OTU where the full-length reference sequence had more than 5 degenerate nucleotides [13]. Pseudo samples were generated by randomly selecting 10 samples with replacement, filtering features to retain only those present in at least 4 samples, averaging the abundance across all samples, and retaining features present with a relative abundance of at least 1/5000 sequences. Simulated samples were scaled to 50,000 sequences.

QIIME 2 feature classifier (q2-feature-classifier; v 2020.11) was used to extract 16S hypervariable regions for each of the primer pairs from the Greengenes 13_8 database’s 97% representative sequences using default parameters [1,2,13]. We expanded all degenerate bases in the database regions, giving equal weight to each degenerate variant. We assumed samples were denoised after sequencing and denoising was perfect. We also assumed the best available short read technology could consistently give us 2x250nt reads after barcode removal and trimming. (Illumina does offer a 2x300 kit, however, primer and barcode removal may still be necessary, as well as quality-trimming.) This was insufficient to span the width of the V13 region, so we used only a forward read for the V13 region. Based on the expected amplicon length, a paired end read would be possible for the V68 region, and so this was trimmed to 400nt.

*Real Vaginal Data Processing*

Vaginal samples were acquired from ENA. For this analysis, we used the samples amplified using a two-step amplification protocol with 27F-534R (V13) and 341-805 (V34) primers (Table S1). The data was denoised using DADA2 in q2-dada2 using default parameters [5]. Single-end denoising was used on the V13 region trimmed to 280nt; single-end reads were trimmed to a final length of 275nt using q2-sidle. The V34 region was denoised using a paired-end strategy with both forward and reverse reads trimmed to 275nt; the final ASVs were trimmed to 350nt for uniformity.

**Table S1. Regional Primers and trim length**

| **Region** | **Ref** | | **Forward** | **Reverse** | **Amplicon Length** |
| --- | --- | --- | --- | --- | --- |
| Simulation Primers | | | | | |
| V1-3  (27F-534R) | [14] | AGAGTTTGATCCTGGCTCAG | | ATTACCGCGGCTGCTGG | 250 |
| V4  (515F-806R) | [15] | GTGYCAGCMGCCGCGGTAA | | GGACTACNVGGGTWTCTAAT | 250 |
| V6-8  (969F-1406R) | [16] | ACGCGHNRAACCTTACC | | ACGGGCRGTGWGTRCAA | 400 |
| Multiple Region Vaginal Pool Primers | | | | | |
| V1-3  (27F-534R) | [14,17] | AGAGTTTGATCCTGGCTCAG | | ATTACCGCGGCTGCTGG | 275 |
| V3-4  (341F-805R) | [17] | CCTACGGGNGGCWGCAG | | GACTACHVGGGTATCTAATCC | 350 |
| SMURF Benchmarking Data | | | | | |
| 74F-315R | [8] | TGGCGGACGGGTGAGTAA | | CTGCTGCCTCCCGTAGGA | 215 |
| 316F-484R | [8] | TCCTACGGGAGGCAGCAG | | TATTACCGCGGCTGCTGG | 125 |
| 486F-650R | [8] | CAGCAGCCGCGGTAATAC | | CGCATTTCACCGCTACAC | 125 |
| 752F-911R | [8] | AGGATTAGATACCCTGGT | | GAATTAAACCACATGCTC | 125 |
| 901F-1057R | [8] | GCACAAGCGGTGGAGCAT | | CGCTCGTTGCGGGACTTA | 125 |
| 1143F-1336R | [8] | AGGAAGGTGGGGATGACG | | CCCGGGAACGTATTCACC | 175 |

**Table S2. Runtime comparisons for SMURF and Sidle**

|  | | | **Total Run time**  **(hh:mm:ss)** | | **Mean**^c^ **runtime per sample**  **(mm:ss)** | | **Mean (std) run time per region**  **(mm:ss)** | |
| --- | --- | --- | --- | --- | --- | --- | --- | --- |
|  |  |  | **SMURF** | **Sidle** | **SMURF** | **Sidle** | **SMURF** | **Sidle** |
| Tutorial Data, Greengenes | | | 01:50:46 | 01:02:49 | 01:50:46 | 01:02:49 |  |  |
|  | Greengenes 6 regions | | 01:43:48 ^b^ | 00:27:03 | N/A | N/A |  |  |
|  |  | Database filtering | --^b^ | 00:02:09 | N/A | N/A | --^b^ | N/A |
|  |  | Regional database preparation | 01:43:48 | 00:24:54 | N/A | N/A | 17:18c | 04:09 (00:36) |
|  | Sample Preparation | | 00:07:56 | 00:35:46 |  |  |  |  |
|  |  | Regional denoising/quality filtering | 00:00:41 | 00:12:45 | 00:41 | 00:12:45 | 00:07^c^ | 02:13 (00:38) |
|  |  | Alignment | 00:04:35 | 00:07:02 | 04:35 | 00:07:02 | 00:46^c^ | 01:10 (00:24) |
|  |  | Reconstruction | 00:01:42 | 00:15:59 | 01:42 | 00:15:53 | N/A | N/A |
| Vaginal Data,^a^ Greengenes | | |  | 00:44:35 |  | 00:01:55 |  |  |
|  | Greengenes 2 regions | | 01:26:18 ^b^ | 00:14:48 | N/A | N/A |  |  |
|  |  | Database filtering | --^b^ | 00:01:19 | N/A | N/A | --^b^ | N/A |
|  |  | Regional database preparation | 01:26:18 | 00:13:29 | N/A | N/A | 41:54^c^ | 06:44 (00:35) |
|  | Sample Preparation, Greengenes | | --^d^ | 00:29:47 | --^d^ | 00:01:14 | --^d^ |  |
|  |  | Regional denoising/quality filtering | --^d^ | 00:17:26 | --^d^ | 00:00:44 | --^d^ | 08:43 (03:06) |
|  |  | Alignment | --^d^ | 00:13:29 | --^d^ | 00:00:30 | --^d^ | 05:59 (00:37) |
|  |  | Table reconstruction | --^d^ | 00:00:17 | --^d^ | 00:00:01 | --^d^ | N/A |
|  |  | Taxonomic reconstruction | --^d^ | 00:00:06 | --^d^ | N/A | --^d^ | N/A |
| Vaginal Data, Optivag^a,b^ | | |  | 00:21:25 |  | 00:00:54 ^c^ |  |  |
|  | Database Preparation | | --^b^ | 00:01:17 | --^b^ | N/A | --^b^ | 00:39 (00:34) |
|  |  | Regional database preparation | --^b^ | 00:01:17 | --^b^ | N/A | --^b^ | 00:39 (00:34) |
|  | Sample Preparation | | --^b^ | 00:20:08 | --^b^ | 00:00:50 | --^b^ |  |
|  |  | Regional denoising/quality filtering | --^b^ | 00:15:14 | --^b^ | 00:00:38 | --^b^ | 07:37 (03:00) |
|  |  | Alignment | --^b^ | 00:04:41 | --^b^ | 00:00:03 | --^b^ | 02:20 (03:00) |
|  |  | Table reconstruction | --^b^ | 00:00:08 | --^b^ | >00:00:01 | --^b^ | N/A |
|  |  | Taxonomic reconstruction | --^b^ | 00:00:05 | --^b^ | N/A | --^b^ | N/A |
| ^a^Based on custom MATLAB script written to profile multiple samples. The SMURF repository did not contain a similar script. | | | | | | | | |
| ^b^The MATLAB implementation was missing a necessary subfunction to allow degenerate filtering and expansion | | | | | | | | |
| ^c^Calculated as the total time divided by the number of samples | | | | | | | | |
| ^d^SMURF database failed to open due to a difference between the sequence and primer IDs | | | | | | | | |

**Table S3.** Comparison of reconstruction methods on class level data

| **Method** | | | **Mean Relative Abundance** | | **Relationship with Reference** | | | | | |
| --- | --- | --- | --- | --- | --- | --- | --- | --- | --- | --- |
|  |  |  | **Infant** | **Adult** | **Slope** | **[95% CI]** | | **R2** | **fdr p-value** | |
| p Actinobacteria | | |  |  |  |  |  |  |  |  |
|  | c__Actinobacteria | |  |  |  |  |  |  |  |  |
|  |  | reference | 0.017 | 0.230 | 1.00 |  |  |  |  |  |
|  |  | otus | 0.016 | 0.219 | 0.96 | 0.95 | 0.96 | 1.000 | 9E-08 |  |
|  |  | asvs | 0.016 | 0.219 | 0.95 | 0.95 | 0.96 | 1.000 | 4E-08 |  |
|  |  | sidle | 0.017 | 0.229 | 1.00 | 1.00 | 1.00 | 1.000 | 1.00 |  |
|  | c__Coriobacteriia | |  |  |  |  |  |  |  |  |
|  |  | reference | 4.2E-03 | 4.6E-03 | 1.00 |  |  |  |  |  |
|  |  | otus | 4.3E-03 | 4.3E-03 | 0.95 | 0.94 | 0.97 | 0.997 | 0.045 |  |
|  |  | asvs | 4.2E-03 | 4.3E-03 | 0.95 | 0.94 | 0.96 | 0.997 | 0.010 | * |
|  |  | sidle | 4.1E-03 | 4.6E-03 | 0.98 | 0.98 | 0.99 | 0.999 | 0.002 |  |
| Firmictues | | |  |  |  |  |  |  |  |  |
|  | c__Clostridia | |  |  |  |  |  |  |  |  |
|  |  | reference | 0.662 | 0.355 | 1.00 |  |  |  |  |  |
|  |  | otus | 0.624 | 0.295 | 0.92 | 0.90 | 0.93 | 0.996 | 1E-21 | * |
|  |  | asvs | 0.620 | 0.278 | 0.90 | 0.89 | 0.92 | 0.993 | 9E-24 | * |
|  |  | sidle | 0.624 | 0.275 | 0.91 | 0.89 | 0.93 | 0.993 | 2E-22 | * |
|  | c__Bacilli | |  |  |  |  |  |  |  |  |
|  |  | reference | 0.010 | 0.108 | 1.00 |  |  |  |  |  |
|  |  | otus | 0.008 | 0.105 | 0.97 | 0.96 | 0.98 | 0.998 | 7E-03 |  |
|  |  | asvs | 0.009 | 0.121 | 1.13 | 1.11 | 1.15 | 0.996 | 6E-03 | * |
|  |  | sidle | 0.009 | 0.115 | 1.08 | 1.06 | 1.09 | 0.996 | 0.26 |  |
|  | c__Erysipelotrichia | |  |  |  |  |  |  |  |  |
|  |  | reference | 0.024 | 0.029 | 1.00 |  |  |  |  |  |
|  |  | otus | 0.033 | 0.032 | 1.14 | 1.10 | 1.18 | 0.982 | 8E-15 | * |
|  |  | asvs | 0.032 | 0.032 | 1.14 | 1.10 | 1.18 | 0.983 | 3E-15 | * |
|  |  | sidle | 0.027 | 0.030 | 1.04 | 1.03 | 1.05 | 0.997 | 8E-09 |  |
| Bacteriodetes | | |  |  |  |  |  |  |  |  |
|  | c__Bacteroidia | |  |  |  |  |  |  |  |  |
|  |  | reference | 0.241 | 0.163 | 1.00 |  |  |  |  |  |
|  |  | otus | 0.246 | 0.161 | 1.01 | 1.00 | 1.02 | 0.999 | 0.60 |  |
|  |  | asvs | 0.250 | 0.162 | 1.02 | 1.02 | 1.03 | 0.999 | 3E-04 |  |
|  |  | sidle | 0.242 | 0.163 | 1.00 | 1.00 | 1.01 | 1.000 | 0.78 |  |

| **Method** | | | **Mean Relative Abundance** | | **Relationship with Reference** | | | | | |
| --- | --- | --- | --- | --- | --- | --- | --- | --- | --- | --- |
|  |  |  | **Infant** | **Adult** | **Slope** | **[95% CI]** | | **R2** | **fdr p-value** | |
| Proteobacteria | | |  |  |  |  |  |  |  |  |
|  | c__Gammaproteobacteria | |  |  |  |  |  |  |  |  |
|  |  | reference | 0.020 | 0.090 | 1.00 |  |  |  |  |  |
|  |  | otus | 0.021 | 0.087 | 0.97 | 0.96 | 0.98 | 0.999 | 0.02 |  |
|  |  | asvs | 0.020 | 0.088 | 0.98 | 0.97 | 0.98 | 0.999 | 0.24 |  |
|  |  | sidle | 0.018 | 0.088 | 0.97 | 0.96 | 0.99 | 0.997 | 4E-04 |  |
|  | c__Betaproteobacteria | |  |  |  |  |  |  |  |  |
|  |  | reference | 7.4E-03 | 6.5E-03 | 1.00 |  |  |  |  |  |
|  |  | otus | 7.4E-03 | 6.4E-03 | 0.99 | 0.97 | 1.00 | 0.996 | 1.00 |  |
|  |  | asvs | 7.3E-03 | 6.4E-03 | 0.98 | 0.96 | 0.99 | 0.996 | 1.00 |  |
|  |  | sidle | 7.5E-03 | 6.5E-03 | 1.01 | 1.01 | 1.01 | 1.000 | 9E-05 |  |
|  | c__Deltaproteobacteria | |  |  |  |  |  |  |  |  |
|  |  | reference | 3.1E-03 | 7.3E-04 | 1.00 |  |  |  |  |  |
|  |  | otus | 3.7E-03 | 8.1E-04 | 1.19 | 1.16 | 1.21 | 0.993 | 2E-05 | * |
|  |  | asvs | 3.7E-03 | 8.1E-04 | 1.18 | 1.16 | 1.20 | 0.994 | 2E-05 | * |
|  |  | sidle | 3.3E-03 | 7.4E-04 | 1.03 | 1.03 | 1.04 | 1.000 | 2E-05 |  |
|  | c__Epsilonproteobacteria | |  |  |  |  |  |  |  |  |
|  |  | reference | 6.4E-05 | 6.0E-03 |  |  |  |  |  |  |
|  |  | otus | 6.0E-05 | 5.6E-03 | 0.93 | 0.92 | 0.94 | 0.999 | 0.005 | * |
|  |  | asvs | 5.9E-05 | 5.6E-03 | 0.93 | 0.92 | 0.94 | 0.999 | 0.004 | * |
|  |  | sidle | 6.6E-05 | 6.0E-03 | 1.00 | 1.00 | 1.01 | 1.000 | 1.000 |  |
| Tenerictues | | |  |  |  |  |  |  |  |  |
|  | c__Mollicutes | |  |  |  |  |  |  |  |  |
|  |  | reference | 3.1E-03 | 1.0E-04 | 1.00 |  |  |  |  |  |
|  |  | otus | 3.7E-03 | 9.9E-05 | 1.22 | 1.19 | 1.25 | 0.991 | 0.002 | * |
|  |  | asvs | 3.8E-03 | 9.9E-05 | 1.22 | 1.19 | 1.25 | 0.990 | 0.001 | * |
|  |  | sidle | 3.4E-03 | 9.6E-05 | 1.08 | 1.06 | 1.11 | 0.994 | 0.030 | * |
| Verrumicrobia | | |  |  |  |  |  |  |  |  |
|  | c__Verrucomicrobiae | |  |  |  |  |  |  |  |  |
|  |  | reference | 6.4E-03 | 5.5E-03 | 1.00 |  |  |  |  |  |
|  |  | otus | 6.1E-03 | 5.3E-03 | 0.95 | 0.94 | 0.96 | 0.998 | 0.018 |  |
|  |  | asvs | 6.1E-03 | 5.3E-03 | 0.95 | 0.94 | 0.96 | 0.998 | 0.005 | * |
|  |  | sidle | 6.6E-03 | 5.5E-03 | 1.01 | 1.01 | 1.01 | 1.000 | 0.032 |  |

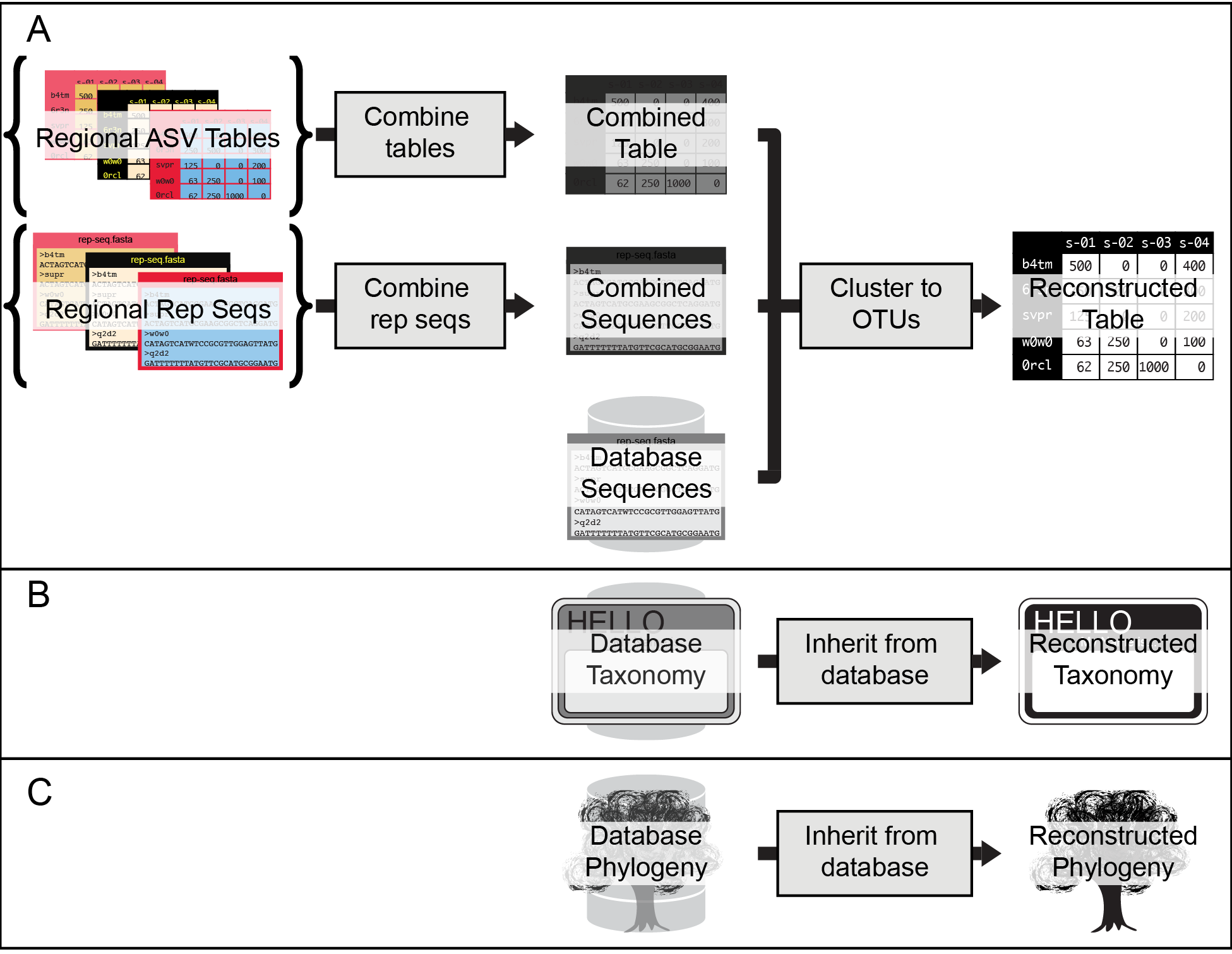

**Figure S1. OTU processing workflow.** Pre-region files and steps are shown in curly brackets ({}). Final reconstructed artifacts are black and white. (A) The ASV tables from all regions and representative sequences from all regions (example region shown here in red and gold) are combined, to generate a combined table and combined sequences, respectively (Dark gray). These are clustered into OTUs using the reference database. The (B) taxonomy and (C) phylogeny come directly from the database.

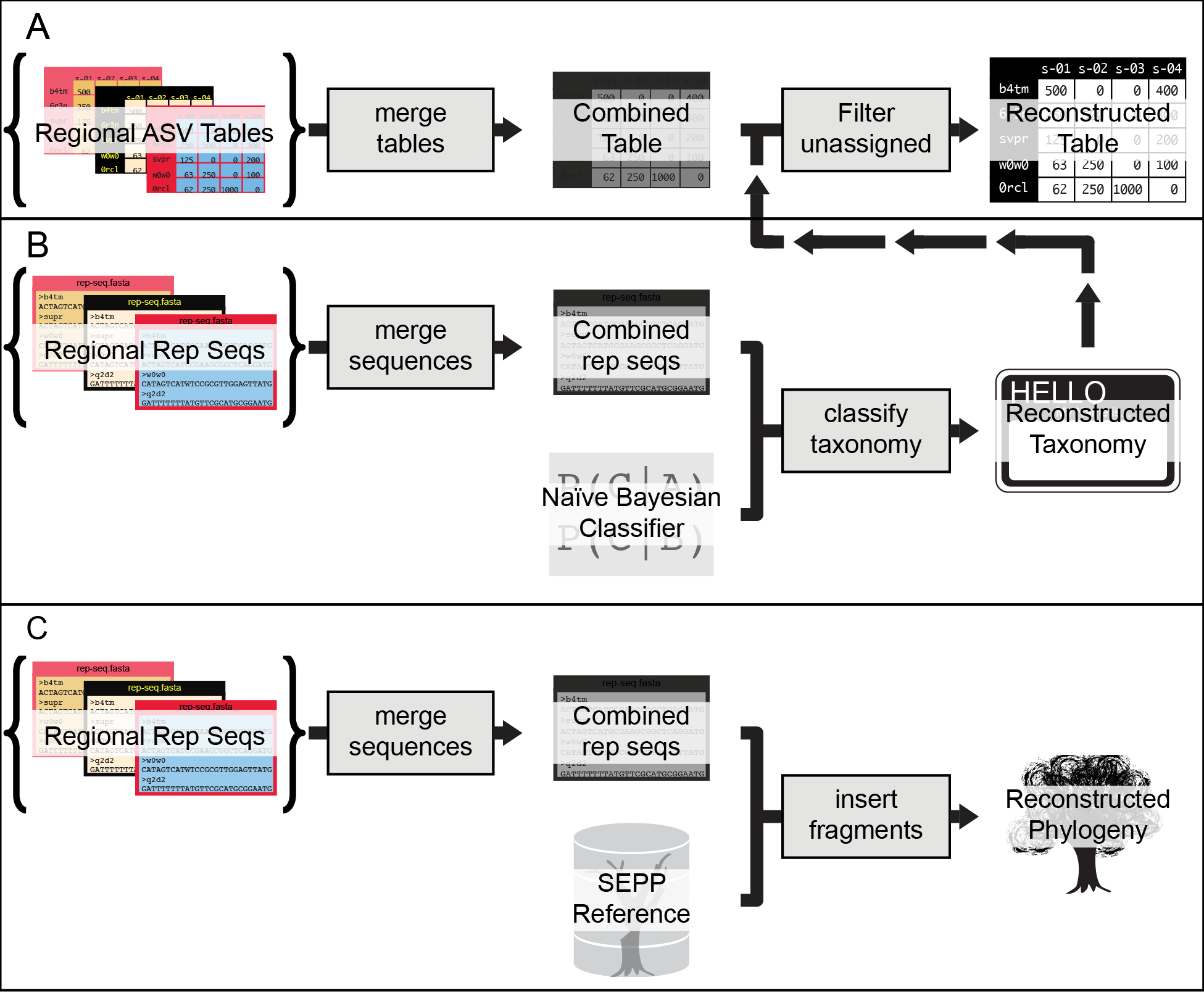

**Figure S2. ASV processing workflow** Pre-region files and steps are shown in curly brackets ({}) and colored red and gold. Final reconstructed artifaccts are black and white. (A) Per-region ASV tables from all regions are merged combined to generate a combined table. The final table is filtered to exclude any sequence missing a phylum level designation in the reconstructed taxonomy. (B) Taxonomy is reconstructed by merging all sequences and classifying with a naïve Bayesian classifier trained against the full length 16S rRNA gene sequence. (C) A phylogenetic tree is built by combining all sequences and inserting them into a reference backbone.
